# GUIFold - A graphical user interface for local AlphaFold2

**DOI:** 10.1101/2023.01.19.521406

**Authors:** Georg Kempf, Simone Cavadini

**Affiliations:** Friedrich Miescher Institute for Biomedical Research, Maulbeerstrasse 66, 4058 Basel, Switzerland

**Keywords:** guifold, alphafold, gui, graphical, prediction

## Abstract

GUIFold is a graphical user interface for the open-source structure prediction pipeline AlphaFold2. A particular emphasis lies on tracking prediction jobs in a database as well as user-friendly submission to queueing systems. GUIFold is built on top of a modified AlphaFold2 pipeline that primarily allows more control of the feature generation step. Additionally, the application provides an evaluation pipeline to rank predictions and visualize confidence metrics. GUIFold can be installed along with AlphaFold2 in a virtual environment and, after initial setup, allows running jobs without particular technical expertise.

## Introduction

The AlphaFold Database provides access to millions of precomputed predictions of monomeric protein structures (Varadi et al., 2022). The number of possible protein interactions is several orders of magnitude higher than the size of single target proteomes and prediction of multimeric complexes usually goes hand in hand with large sequence sizes. Such prediction jobs require high-end hardware resources, making it challenging to build-up a comprehensive collection of precalculated complex structures. Thus, along with other specialized applications such as refinement of experimentally derived structure models, there is still a significant demand to run custom prediction jobs. DeepMind provides the complete open-source code to run the AlphaFold2 pipeline from the command line as well as a Google Colaboratory (Colab) notebook for cloud-based custom predictions (Jumper et al., 2021; Evans et al., 2021). In addition, other groups provide Colab notebooks, extending the AlphaFold pipeline by additional features (Mirdita et al., 2021). However, free-of-charge cloud resources are limited and, if appropriate hardware is available, using a locally installed AlphaFold2 appears to us the method of choice. GUIFold was initially developed to facilitate running predictions on our local compute environment. The main features of this graphical user interface (GUI) application comprise a customisable queue dispatcher with automatic resource selection, tracking of jobs in a database, and evaluation of results. The GUI uses a modified version of AlphaFold2 allowing more control of the feature generation pipeline. We made the application publicly available in the hope that others might find it useful.

## Methods

### Implementation

The code basis was adapted from (Kempf, 2021) developed by the same authors. GUIFold is being developed using *Python 3.8* in combination with the *PyQt5* framework. This allows seamless integration into a local environment (e.g. *conda*) along with AlphaFold2 and dependencies. To store information, a *SQLite* database file is created in the home directory of each user. The *SQLAlchemy* package (Bayer, 2010) is used for most database operations. The *Jinja2* package is employed to manage templates for cluster submission scripts and report generation. Report templates are written with HTML and JavaScript. The evaluation pipeline uses the *Matplotlib, NumPy* and *biopython* (Cock et al., 2009) packages. Models are visualized with 3Dmol.js (Rego and Koes, 2015) and PyMOL (Schrödinger, LLC, 2021). Loosely following various GUI design patterns, the GUI functions are divided into several classes to manage separate functionalities such as job parameters, job control, projects, and general settings. The database model is reflected from the main classes and automatically generated. This allows easy extension of the GUI by additional parameters.

### Structure prediction

Predictions were performed with a modified version version of AlphaFold v2.3.0 (https://github.com/fmi-basel/GUIFold, commit e19b77f). Root mean square deviations (r.m.s.d.) between structures were calculated with PyMOL using *cealign*.

## Results

### User Interface

The graphical user interface (Fig. 1) is organized in two parts: 1) on the left side, a notebook with job configuration, job monitoring, and evaluation tabs and 2) on the right side, a project and job management area. As a first step to run a new job, the user pastes a single, or in case of batch or multimer prediction, multiple sequences into the sequence text field (FASTA sequence format required) and reads-in the sequences with a button below. The individual components are then listed in the below “component table”, where for each sequence/component precomputed MSAs (e.g. from a previous run) and/or a custom template (PDBx/mmCIF format required) can be provided. In addition, search for templates and/or MSA generation can be disabled. The path to a collection of precomputed MSAs (e.g. from a previous “multimer” or “batch_msas” job) can be set in the “Precomputed MSAs path” input field. Relaxation of predicted models can be toggled (“Run Relax” option) and different pipeline protocols can be selected from the “Pipeline” list of options (“only_features”, “continue_-from_features”, “batch_msas”). The database presets (“DB preset” list of options) can be switched from “full” to “reduced_dbs” or “colabfold”. In the “Prediction” list of options, the implementation can be switched from default “alphafold” to “fastfold”, which is a performance optimized reimplementation of AlphaFold2 (Cheng et al., 2022). A more detailed explanation of these options and protocols is provided below. Jobs can be run directly on the host or submitted to a queueing system (“Submit to Queue” option). Additional settings from the AlphaFold2 and FastFold pipelines can be accessed in a separate dialog (“Advanced Settings”). After starting a job, the page is automatically switched to the job monitoring page, where the job progression and output from the queueing system and AlphaFold pipeline is displayed. After completion of the job, the page is switched to the Evaluation tab, where the results from the evaluation pipeline are displayed.

**Figure 1.**
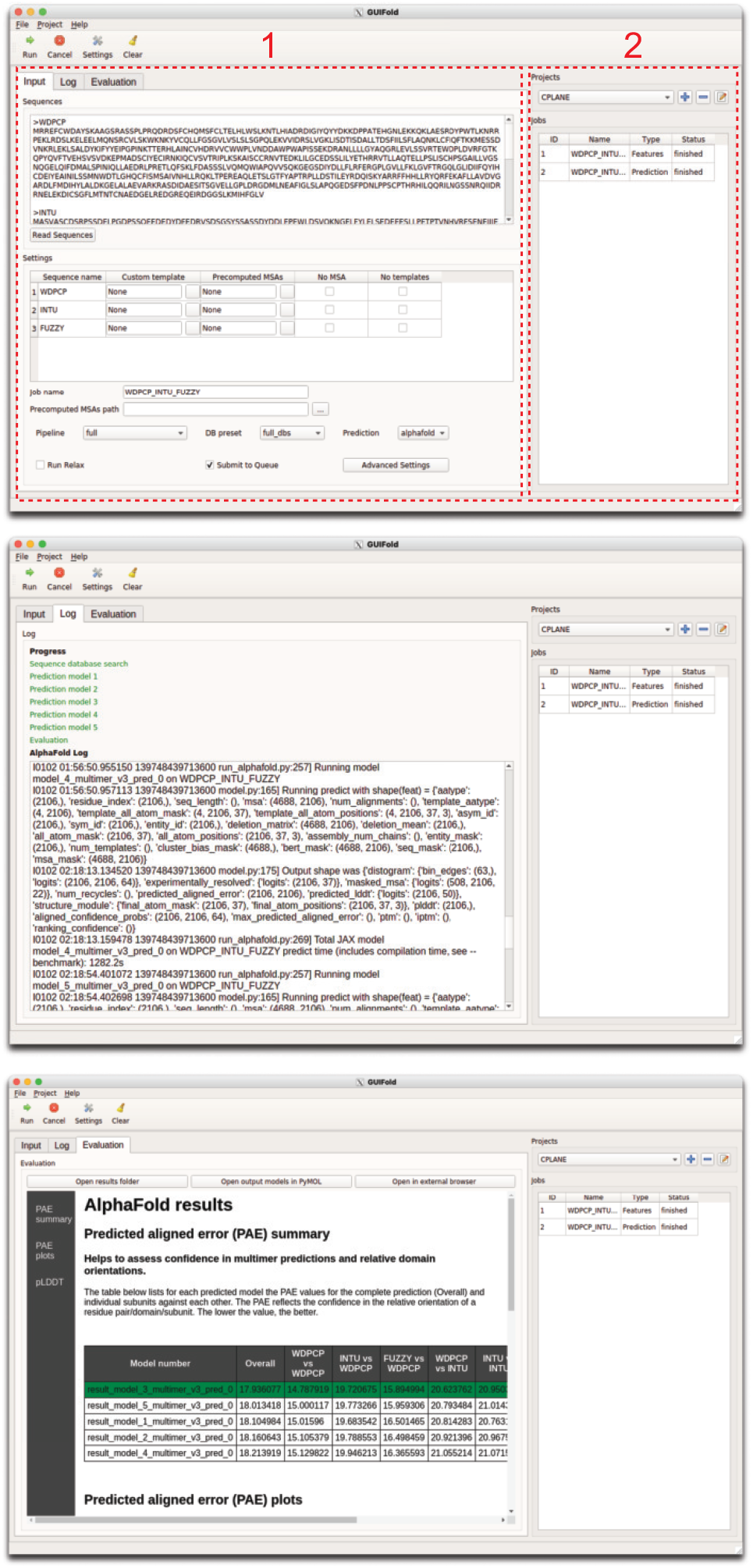
Organization of the GUI. Top panel, Input tab for entering sequences and adjusting settings. (1) Notebook with job configuration/log/evaluation tabs, (2) project and job management area. Middle panel, Log tab showing output from the AlphaFold pipeline and progress. Bottom panel, Evaluation tab displaying confidence metrics (PAE, pLDDT).

### Features

#### Modified feature generation pipeline

GUIFold is built on top of a modified version of the AlphaFold2 pipeline to include support for selective use of precomputed MSAs, a (single) custom template, disabling of MSA generation and/or template search. In addition to the default MSA pipelines (“full” and “reduced_dbs”, see (Jumper et al., 2021)), MSAs can also be generated using MMseqs2 in combination with the uniref30 and colabfold_envdb databases (“colabfold” option) (Steinegger and Söding, 2017). In case of multimer predictions, sequences are still paired through the default protocol of the alphafold pipeline. The workflow to run MMseqs2 was reimplemented based on (Mirdita et al., 2021). When a database index is created and loaded into memory, runtime for MSA generation with MMseqs2 can be significantly reduced (Mirdita et al., 2021).

The whole pipeline (feature generation and prediction) can be split into CPU- and GPU-parts and the multi-threaded database searches are additionally parallelized over two processes (so that HHblits runs in parallel to the faster jackhmmer jobs). To split the job into CPU and GPU parts (e.g. if GPU resources are limited on a compute cluster), the job can be stopped after feature generation (“only_features” option), and then re-submitted using the “continue_from_features” option. There is also an option (“Split job” in settings) to automatically submit two inter-dependent jobs to the queuing system. The second job (CPU/GPU, prediction step) waits until the first job (CPU, feature step) completes. This requires support of job dependencies in the queueing system and the documentation provides an example for SLURM.

A collection of precomputed MSAs can be generated in batch mode when multiple sequences are provided and the option “batch_msas” is selected. In contrast to the multimer pipeline, MSAs will not be paired at this stage and the generated MSAs and template hits can be reused in different combinations in subsequent multimer jobs (where the respective pairing is done). When the path to a job or project folder is provided in the “Precom-puted MSAs path” field, MSA files (from the AlphaFold2 pipeline) will be automatically searched and selected by comparison to the input sequences. If a subsequence is provided, precomputed MSAs will be cropped accordingly. It should be noted that the cropped MSAs might be different from MSAs recalculated for the subsequence and thus might yield different prediction results. When a partially finished previous job is started again with the same “job name” MSAs can be re-used or overwritten (corresponds to “use_precomputed_msas” option of the default AlphaFold2 pipeline). A flowchart of the pipeline control options is provided in Fig. 2.

**Figure 2.**
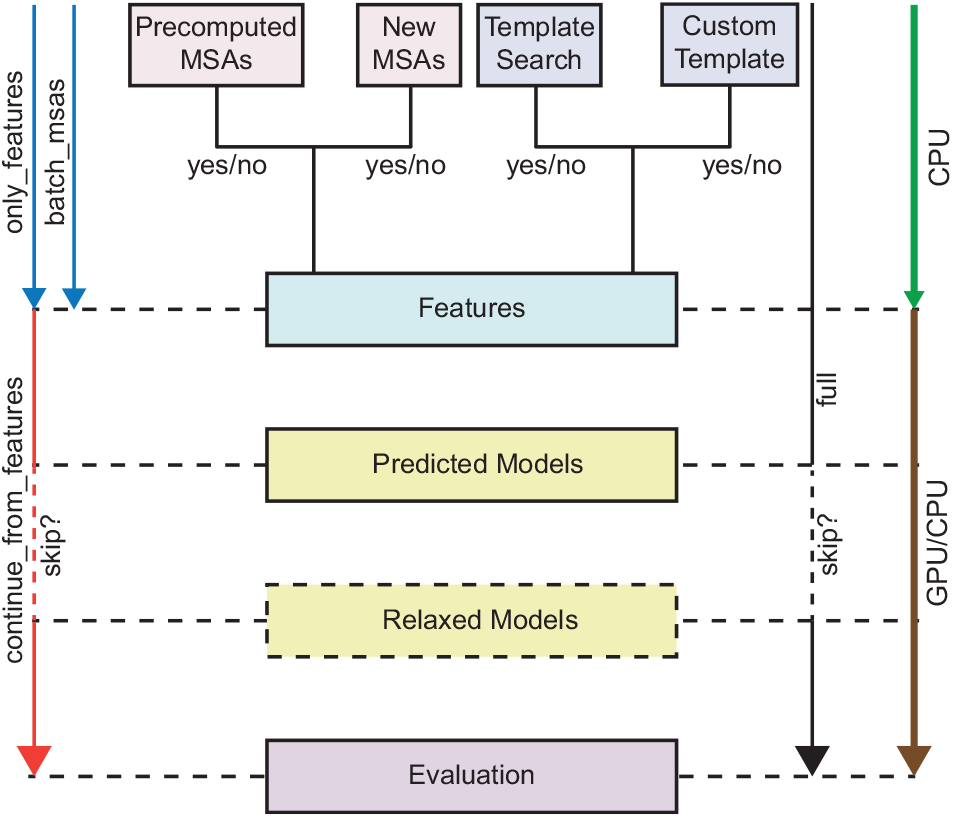
Pipeline control options. The pipeline is separated into feature generation, prediction, model relaxation, and evaluation. Full features are mainly computed from MSAs and templates, which can be selectively disabled. Instead of generating new MSAs, precomputed MSAs can be supplied. A custom template can be used instead of templates from the PDB. The pipeline can be split up allowing to run steps that require only CPU on different resources.

#### Evaluation pipeline

An additional evaluation pipeline was developed that summarizes the pLDDT or PAE values (definition in (Jumper et al., 2021)) and generates plots for the PAE. The PAE between subunits is useful to assess the confidence for a multimer prediction. Therefore, in a table, the overall PAEs and PAEs extracted for all possible combinations of subunits are listed for each model. This report is saved as a HTML file in the results folder from the AlphaFold2 pipeline and can be displayed in the results tab of the GUI or opened in an external browser. An additional HTML file is generated that includes the 3Dmol.js viewer (Rego and Koes, 2015) to visualize the predicted models (Supp. Fig. 3). The user can switch between coloring by pLDDT and different chains. A button to open the output structure models in PyMOL is provided in the evaluation page.

#### Automation of submission to queuing systems

System administrators can configure the settings (e.g. genetic database locations, queuing system) in a configuration file that will be automatically used by other instances when the software is installed in a shared storage. Cluster resource selection can be controlled through required and optional variables in the submission template (Tab. 1). Based on the total sequence length, the required GPU memory is estimated and can be accessed through a variable. If multiple GPUs with different memory are available, the estimated memory can be used to select the appropriate GPU in the submission script. Additionally, it is possible to control the use of unified memory in case the GPU memory would be exceeded. An example submission template for the SLURM queuing system is provided along with the source code in the “templates” folder. The submission template only needs to be configured once and will then be used by all other instances started from a shared installation location.

**Table 1.** Available submission template variables. An example script, illustrating the use of these variables, is provided in the “templates” folderofthe source code.

| Template variable | Description |
| --- | --- |
| {{logfile}} | (required) Path to the log file. |
| {{account}} | (optional) When a specific account is needed to run jobs on the cluster. |
| {{use_gpu}} | (optional) To build a conditional block for selecting CPU or GPU resources. The variable will hold False if no GPU resources are required for the job type (e.g. "Only Features" or "Force CPU"). |
| {{mem}} | (optional) How much RAM (in GB) should be reserved. The RAM will be automatically increased with the GPU memory in case of unified memory (up to the defined maximum RAM). |
| {{num_cpu}} | (optional) Number of CPUs to request. |
| {{num_gpu}} | (optional) Number of GPUs to request (only relevant when FastFold implementation is used). |
| {{total_sequence_length}} | (optional) Total sequence length (including identical sequences). |
| {{gpu_mem}} | (optional) Useful when the queuing system supports selection of GPU by memory requirement. Value in GB. |
| {{split_mem}} | (optional) If the required memory exceeds the available GPU memory, the job can be run with unified memory. The split_mem variable holds None or the memory split fraction and can be used for a conditional block to set the flags required to enable unified memory use. |
| {{add_dependency}} | (optional) When the job is started with "split_job setting", this variable will be True for the second job (prediction step) and allows adding a dependency on the first job (feature step). |
| {{queue_job_id}} | (required) Job ID from the queuing system. |
| {{command}} | (required) The command to run the AlphaFold job. |

#### Interface to FastFold

As part of the OpenFold project, AlphaFold2 has been reimplemented using the PyTorch framework (Ahdritz et al., 2022). The FastFold project builds on OpenFold and offers additional performance optimizations such as lower GPU memory consumption and faster runtimes (Cheng et al., 2022). GUIFold provides the option to use FastFold for the prediction step (starting from the feature processing step). When FastFold is selected for prediction, calculation of MSAs and template search is still carried out with the above described modified AlphaFold2 pipeline. Under “Advanced Settings”, FastFold specific parameters can be adjusted. The installation script automatically installs FastFold and dependencies in the same environment as GUIFold and AlphaFold2.

#### Other features

Under “Advanced Settings” the number of model recycling steps can be adjusted and a higher number has been shown to improve the prediction quality in difficult cases (Mirdita et al., 2021). The application supports launching and monitoring multiple jobs and the user can create projects to organize related predictions, for example when several jobs are to be run in the context of a particular research project. When a previous job is selected, the settings and results for this job are loaded into the respective notebook pages. This also allows to start a modified job without having to setup all parameters from scratch.

#### Related work

Some of the described features are available in a similar manner in other packages and were implemented independently for this project (Mirdita et al., 2021; Zhong et al., 2022; Yu et al., 2022).

### Usage example

The utility of GUIFold was demonstrated by predicting the human CPLANE core complex (usage example in supplementary information). The r.m.s.d between the predicted and experimental structure (Langousis et al., 2022) was 1.5 Å. In another example, binding of the small GTPase RSG1 to the CPLANE core complex is predicted. In this case, the MSAs from the previous job (CPLANE core complex) were re-used. The predicted position of RSG1 is in good agreement with the experimental structure of the mouse CPLANE-RSG1 complex (1.9 Å r.m.s.d between predicted human and experimental mouse (Langousis et al., 2022) CPLANE-RSG1 structures).

## Discussion and Outlook

GUIFold provides a graphical user interface for a modified version of the open-source AlphaFold2 pipeline. It is specifically intended for the use on multi-user systems and compute clusters.

Although the AlphaFold database provides millions of precalculated predictions, a significant need for calculating custom predictions remains. Among various specialized applications, prediction of multimeric protein assemblies constitutes the most significant task. When suitable hardware infrastructure is available, running AlphaFold2 locally appears to us the method of choice. GUI-based applications help to make this option better accessible.

## Software Availability

The source code of GUIFold is available from https://github.com/fmi-basel/GUIFold and is licensed under the Apache 2.0 License. Installation instructions and further documentation are provided at https://github.com/fmi-basel/GUIFold#readme.

## Acknowledgments

We thank David Domjan, Jakob Schnabl, Fabio Mohn and Si Hoon Park for testing the software. This work was supported by the Novartis Research Foundation.

