## Supplementary Information for "GUIFold - A graphical user interface for local AlphaFold2"

### 1 Usage example

The below described example assumes that GUIFold is already installed and configured, the application is launched, and a project has been created (see documentation https://github.com/fmi-basel/GUIFold/blob/main/README.md for further guidelines).

#### 1.1 Input sequences

Uniprot entries: O95876 (WDPCP), Q9ULD6 (INTU), Q9BT04 (FUZZY)

>WDPCP
MRREFCWDAYSKAAGSRASSPLPRQDRDSFCHQMSFCLTELHLWSLKNTLHIADRDIGIYQYYDKKDPPATEHGNLEKKQKLAESRDYPWTLKNRRPEKLRDSLKELEELMQNSRCVLSKWKNKYVCQLLFGS
GVLVSLSLSGPQLEKVVIDRSLVGKLISDTISDALLTDSFIILSFLAQNKLCFIQFTKKMESSDVNKRLEKLSALDYKIFYYEIPGPINKTTERHLAINCVHDRVVCWWPLVNDDAWPWAPISSEKDRANLLLLGYAQGR
LEVLSSVRTEWDPLDVRFGTKQPYQVFTVEHSVSVDKEPMADSCIYECIRNKIQCVSVTRIPLKSKAISCCRNVTEDKLILGCEDSSLILYETHRRVTLLAQTELLPSLISCHPSGAILLVGSNQGELQIFDMALSPINI
QLLAEDRLPRETLQFSKLFDASSSLVQMQWIAPQVVSQKGEGSDIYDLLFLRFERGPLGVLLFKLGVFTRGQLGLIDIIFQYIHCDEIYEAINILSSMNWDTLGHQCFISMSAIVNHLLRQKLTPEREAQLETSLGTFY
APTRPLLDSTILEYRDQISKYARRFFHHLLRYQRFEKAFLLAVDVGARDLFMDIHYLALDKGELALAEVARKRASDIDAESITSGVELLGPLDRGDMLNEAFIGLSLAPQGEDSFPDNLPPSCPTHRHILQQRILNGSS
NRQIIDRRNELEKDICSGFLMTNTCNAEDGELREDGREQEIRDGGSLKMIHFGLV

>INTU
MASVASCDSRPSSDELPGDPSSQEEDEDYDFEDRVSDSGSYSSASSDYDDLEPEWLDSVQKNGELFYLELSEDEEESLLPETPTVNHVRFSENEIIIEDDYKERKKYEPKLKQFTKILRRKRLLPKRCNKKNSND
NGPVSILKHQSNQKTGVIVQQRYKDVNVYVNPKKLTVIKAKEQLKLLEVLVGIIHQTKWSWRRTGKQGDGERLVVHGLLPGGSAMKSGQVLIGDVLVAVNDVDVTTENIERVLSCIPGPMQVKLTFENAYDVKRET
SHPRQKKTQSNTSDLVKLLWGEEVEGIQQSGLNTPHIIMYLTLQLDSETSKEEQEILYHYPMSEASQKLKSVRGIFLTLCDMLENVTGTQVTSSSLLLNGKQIHVAYWKESDKLLLIGLPAEEVPLPRLRNMIENVIQ
TLKFMYGSLDSAFCQIENVPRLDHFFNLFFQRALQPAKLHSSASPSAQQYDASSAVLLDNLPGVRWLTLPLEIKMELDMALSDLEAADFAELSEDYYDMRRLYTILGSSLFYKGYLICSHLPKDDLIDIAVYCRHYCL
LPLAAKQRIGQLIIWREVFPQHHLRPLADSSTEVFPEPEGRYFLLVVGLKHYMLCVLLEAGGCASKAIGSPGPDCVYVDQVKTTLHQLDGVDSRIDERLASSPVPCLSCADWFLTGSREKTDSLTTSPILSRLQGTS
KVATSPTCRRTLFGDYSLKTRKPSPSCSSGGSDNGCEGGEDDGFSPHTTPDAVRKQRESQGSDGLEESGTLLKVTKKKSTLPNPFHLGNLKKDLPEKELEIYNTVKLTSGPENTLFHYVALETVQGIFITPTLEEV
AQLSGSIHPQLIKNFHQCCLSIRAVFQQTLVEEKKKGLNSGDHSDSAKSVSSLNPVKEHGVLFECSPGNWTDQKKAPPVMAYWVVGRLFLHPKPQELYVCFHDSVTEIAIEIAFKLFFGLTL

>FUZZY
MGEEGTGGTVHLLCLAASSGVPLFCRSSRGGAPARQQLPFSVIGSLNGVHMFGQNLEVQLSSARTENTTVVWKSFHDSITLIVLSSEVGISELRLERLLQMVFGAMVLLVGLEELTNIRNVERLKKDLRASYCLIDS
FLGDSELIGDLTQCVDCVIPPEGSLLQEALSGFAEAAGTTFVSLVVSGRVVAATEGWWRLGTPEAVLLPWLVGSLPPQTARDYPVYLPHGSPTVPHRLLTLTLLPSLELCLLCGPSPPLSQLYPQLLERWWQPLL
DPLRACLPLGPRALPSGFPLHTDILGLLLLHLELKRCLFTVEPLGDKEPSPEQRRRLLRNFYTLVTSTHFPPEPGPPEKTEDEVYQAQLPRACYLVLGTEEPGTGVRLVALQLGLRRLLLLLSPQSPTHGLRSLATH
TLHALTPLL

#### 1.2 Prediction job (CPLANE core complex)

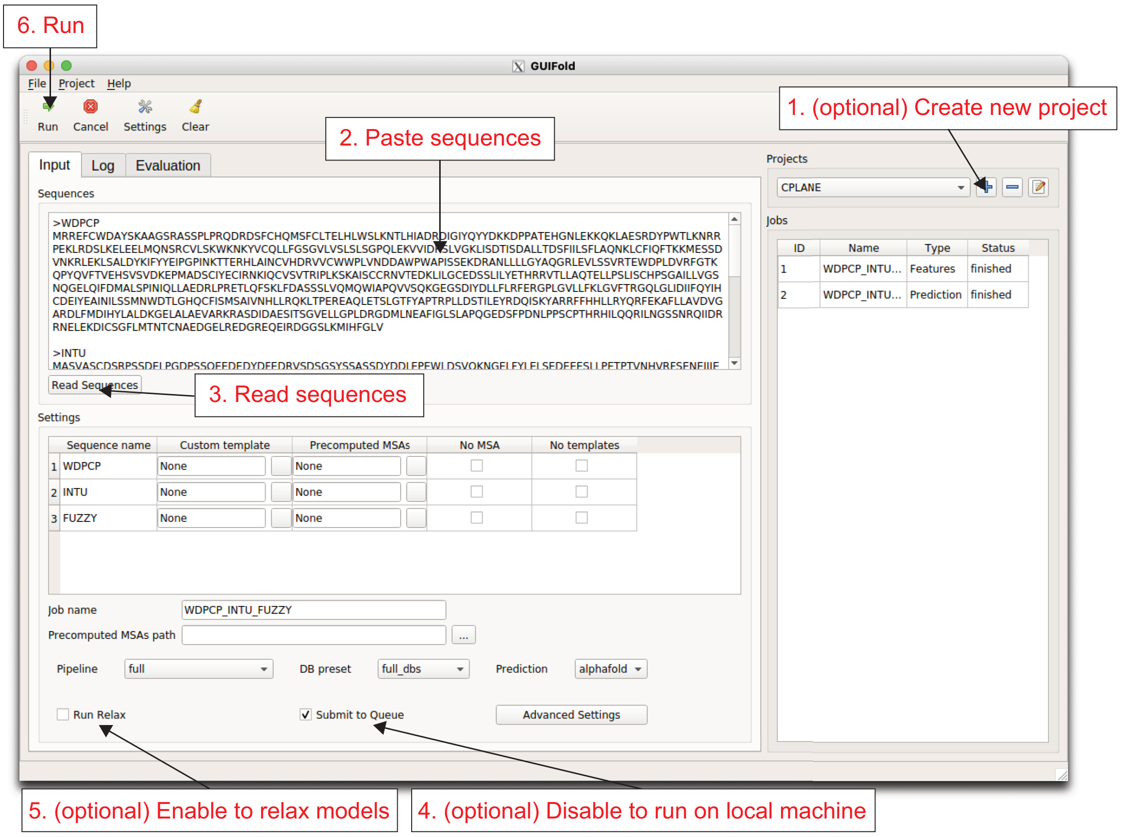


**Supplemental Figure 1: Steps required to start a standard multimer prediction job.**

#### 1.3 Job monitoring and evaluation of predictions

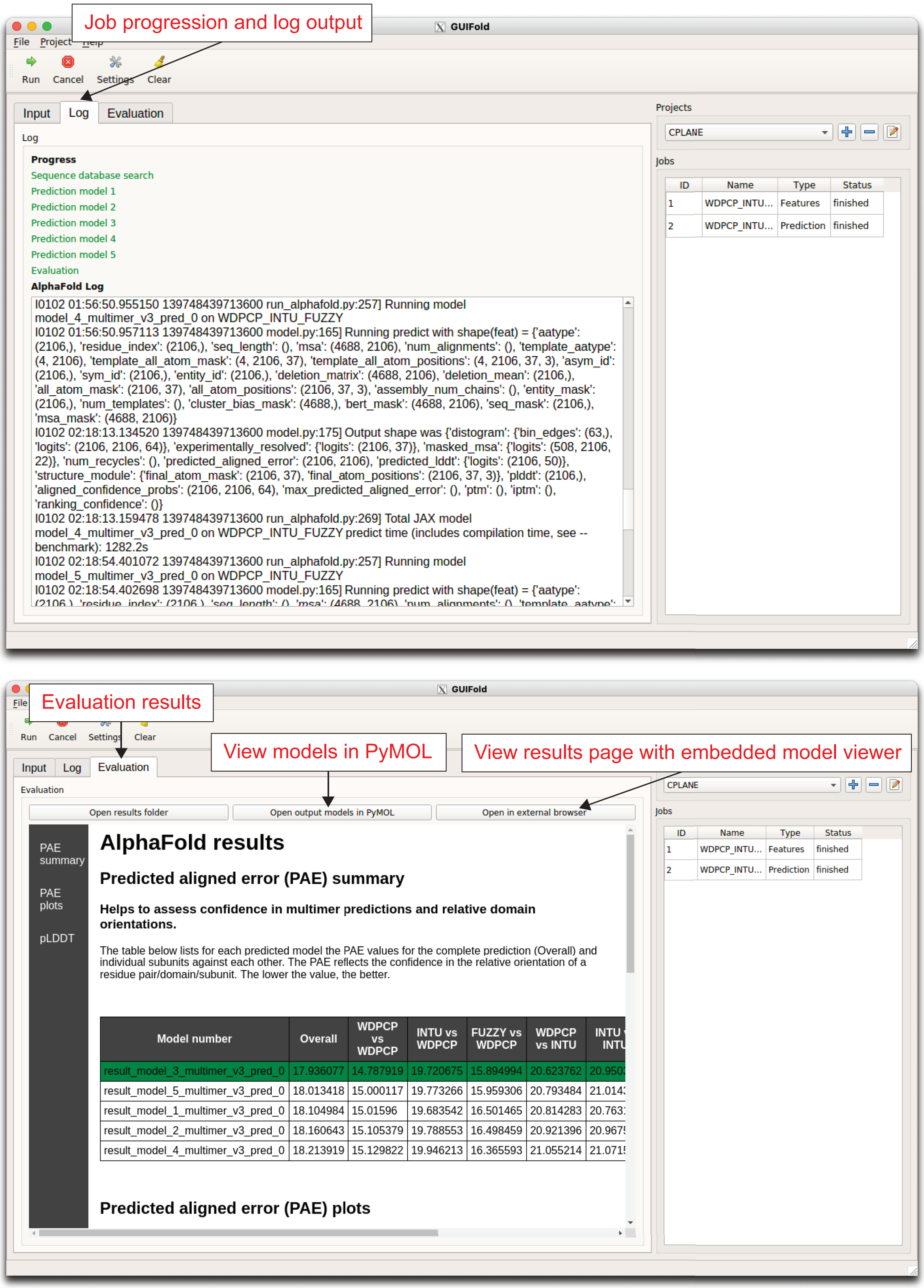


**Supplemental Figure 2: Job monitoring (top) and evaluation (bottom) pages.** To view the models, PyMOL can be launched from the evaluation page and models will be automatically loaded. Two PyMOL scenes are generated where models are colored by pLDDT or chains. Alternatively, an extended results page with embedded model viewer (3Dmol.js) can be opened in an external browser and models can be colored by pLDDT or chains (see below).

#### 1.4 Full evaluation results for CPLANE prediction

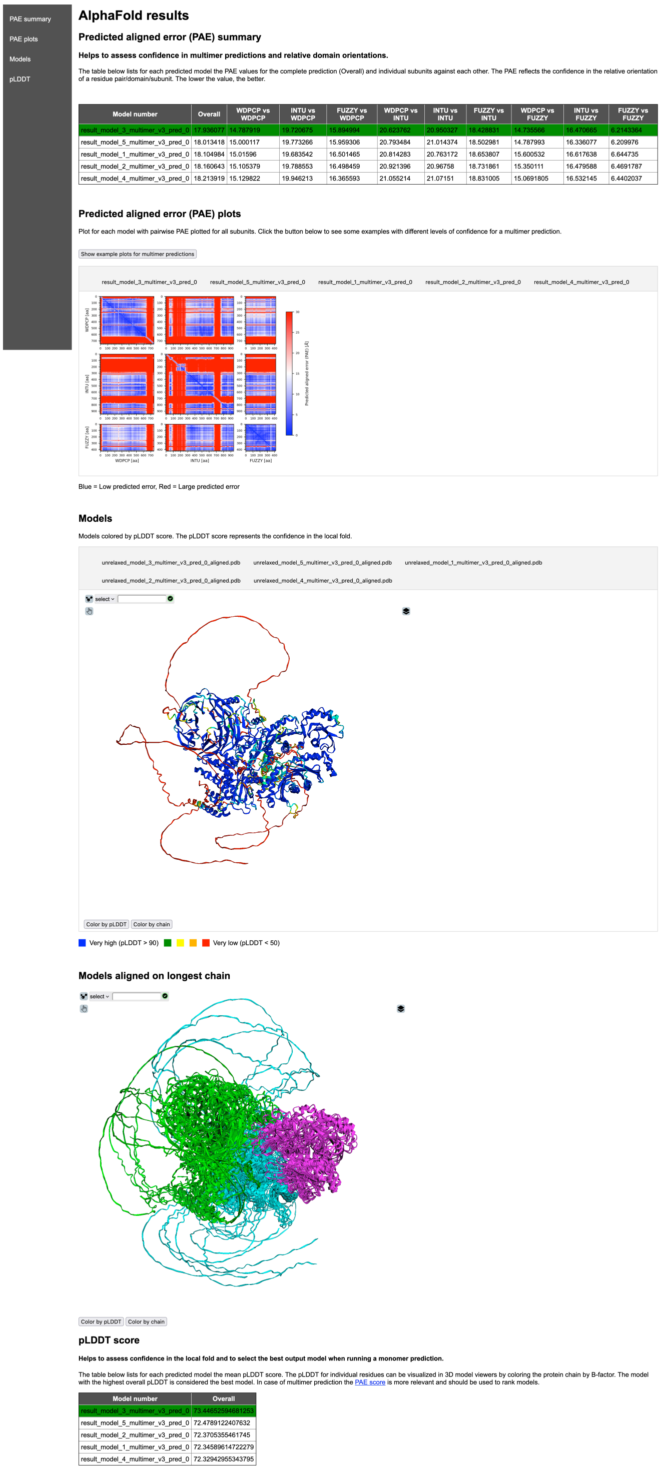

PAE summary
PAE plots
Models
pLDDT

##### AlphaFold results

###### Predicted aligned error (PAE) summary

**Helps to assess confidence in multimer predictions and relative domain orientations.**

The table below lists for each predicted model the PAE values for the complete prediction (Overall) and individual subunits against each other. The PAE reflects the confidence in the relative orientation of a residue pair/domain/subunit. The lower the value, the better.

| Model number | Overall | WDPCP vs WDPCP | INTU vs WDPCP | FUZZY vs WDPCP | WDPCP vs INTU | INTU vs INTU | FUZZY vs INTU | WDPCP vs FUZZY | INTU vs FUZZY | FUZZY vs FUZZY |
|---|---|---|---|---|---|---|---|---|---|---|
| result_model_3_multimer_v3_pred_0 | 17.936077 | 14.787919 | 19.720675 | 15.894994 | 20.623762 | 20.950327 | 18.428831 | 14.735566 | 16.470665 | 6.2143364 |
| result_model_5_multimer_v3_pred_0 | 18.013418 | 15.000117 | 19.773266 | 15.959306 | 20.793484 | 21.014374 | 18.502981 | 14.787993 | 16.336077 | 6.209976 |
| result_model_1_multimer_v3_pred_0 | 18.104984 | 15.01596 | 19.683542 | 16.501465 | 20.814283 | 20.763172 | 18.653807 | 15.600532 | 16.617638 | 6.644735 |
| result_model_2_multimer_v3_pred_0 | 18.160643 | 15.105379 | 19.788553 | 16.498459 | 20.921396 | 20.96758 | 18.731861 | 15.350111 | 16.479588 | 6.4691787 |
| result_model_4_multimer_v3_pred_0 | 18.213919 | 15.129822 | 19.946213 | 16.365593 | 21.055214 | 21.07151 | 18.831005 | 15.0691805 | 16.532145 | 6.4402037 |

###### Predicted aligned error (PAE) plots

Plot for each model with pairwise PAE plotted for all subunits. Click the button below to see some examples with different levels of confidence for a multimer prediction.

Show example plots for multimer predictions

result_model_3_multimer_v3_pred_0 result_model_5_multimer_v3_pred_0 result_model_1_multimer_v3_pred_0 result_model_2_multimer_v3_pred_0 result_model_4_multimer_v3_pred_0

Blue = Low predicted error, Red = Large predicted error

###### Models

Models colored by pLDDT score. The pLDDT score represents the confidence in the local fold.

unrelaxed_model_3_multimer_v3_pred_0_aligned.pdb unrelaxed_model_5_multimer_v3_pred_0_aligned.pdb unrelaxed_model_1_multimer_v3_pred_0_aligned.pdb
unrelaxed_model_2_multimer_v3_pred_0_aligned.pdb unrelaxed_model_4_multimer_v3_pred_0_aligned.pdb

select

Color by pLDDT Color by chain

Very high (pLDDT > 90) Very low (pLDDT < 50)

###### Models aligned on longest chain

select

Color by pLDDT Color by chain

###### pLDDT score

**Helps to assess confidence in the local fold and to select the best output model when running a monomer prediction.**

The table below lists for each predicted model the mean pLDDT score. The pLDDT for individual residues can be visualized in 3D model viewers by coloring the protein chain by B-factor. The model with the highest overall pLDDT is considered the best model. In case of multimer prediction the PAE score is more relevant and should be used to rank models.

| Model number | Overall |
|---|---|
| result_model_3_multimer_v3_pred_0 | 73.44652594681253 |
| result_model_5_multimer_v3_pred_0 | 72.4789122407632 |
| result_model_2_multimer_v3_pred_0 | 72.3705355461745 |
| result_model_1_multimer_v3_pred_0 | 72.34589614722279 |
| result_model_4_multimer_v3_pred_0 | 72.32942955343795 |

**Supplemental Figure 3: Screenshot of evaluation results (CPLANE core complex) with embedded model viewer (3Dmol.js).** The file 'results_model_viewer.html' was opened in an external browser. In the first model viewer, models are viewed separately and in the second model viewer, all models aligned on the longest chain are displayed. In both model viewers, models can be colored by pLDDT or chains.

#### 1.5 Additional prediction job extending from precomputed MSAs (CPLANE-RSG1 complex)

Uniprot entries: O95876 (WDPCP), Q9ULD6 (INTU), Q9BT04 (FUZZY), Q9BU20 (RSG1)

>WDPCP
MRREFCWDAYSKAAGSRASSPLPRQDRDSFCHQMSFCLTELHLWSLKNTLHIADRDIGIYQYYDKKDPPATEHGNLEKKQKLAESRDYPWTLKNRRPEKLRDSLKELEELMQNSRCVLSKWKNKYVCQLLFGS
GVLVSLSLSGPQLEKVVIDRSLVGKLISDTISDALLTDSFIILSFLAQNKLCFIQFTKKMESSDVNKRLEKLSALDYKIFYYEIPGPINKTTERHLAINCVHDRVVCWWPLVNDDAWPWAPISSEKDRANLLLLGYAQGR
LEVLSSVRTEWDPLDVRFGTKQPYQVFTVEHSVSVDKEPMADSCIYECIRNKIQCVSVTRIPLKSKAISCCRNVTEDKLILGCEDSSLILYETHRRVTLLAQTELLPSLISCHPSGAILLVGSNQGELQIFDMALSPINI
QLLAEDRLPRETLQFSKLFDASSSLVQMQWIAPQVVSQKGEGSDIYDLLFLRFERGPLGVLLFKLGVFTRGQLGLIDIIFQYIHCDEIYEAINILSSMNWDTLGHQCFISMSAIVNHLLRQKLTPEREAQLETSLGTFY
APTRPLLDSTILEYRDQISKYARRFFHHLLRYQRFEKAFLLAVDVGARDLFMDIHYLALDKGELALAEVARKRASDIDAESITSGVELLGPLDRGDMLNEAFIGLSLAPQGEDSFPDNLPPSCPTHRHILQQRILNGSS
NRQIIDRRNELEKDICSGFLMTNTCNAEDGELREDGREQEIRDGGSLKMIHFGLV

>INTU
MASVASCDSRPSSDELPGDPSSQEEDEDYDFEDRVSDSGSYSSASSDYDDLEPEWLDSVQKNGELFYLELSEDEEESLLPETPTVNHVRFSENEIIIEDDYKERKKYEPKLKQFTKILRRKRLLPKRCNKKNSND
NGPVSILKHQSNQKTGVIVQQRYKDVNVYVNPKKLTVIKAKEQLKLLEVLVGIIHQTKWSWRRTGKQGDGERLVVHGLLPGGSAMKSGQVLIGDVLVAVNDVDVTTENIERVLSCIPGPMQVKLTFENAYDVKRET
SHPRQKKTQSNTSDLVKLLWGEEVEGIQQSGLNTPHIIMYLTLQLDSETSKEEQEILYHYPMSEASQKLKSVRGIFLTLCDMLENVTGTQVTSSSLLLNGKQIHVAYWKESDKLLLIGLPAEEVPLPRLRNMIENVIQ
TLKFMYGSLDSAFCQIENVPRLDHFFNLFFQRALQPAKLHSSASPSAQQYDASSAVLLDNLPGVRWLTLPLEIKMELDMALSDLEAADFAELSEDYYDMRRLYTILGSSLFYKGYLICSHLPKDDLIDIAVYCRHYCL
LPLAAKQRIGQLIIWREVFPQHHLRPLADSSTEVFPEPEGRYFLLVVGLKHYMLCVLLEAGGCASKAIGSPGPDCVYVDQVKTTLHQLDGVDSRIDERLASSPVPCLSCADWFLTGSREKTDSLTTSPILSRLQGTS
KVATSPTCRRTLFGDYSLKTRKPSPSCSSGGSDNGCEGGEDDGFSPHTTPDAVRKQRESQGSDGLEESGTLLKVTKKKSTLPNPFHLGNLKKDLPEKELEIYNTVKLTSGPENTLFHYVALETVQGIFITPTLEEV
AQLSGSIHPQLIKNFHQCCLSIRAVFQQTLVEEKKKGLNSGDHSDSAKSVSSLNPVKEHGVLFECSPGNWTDQKKAPPVMAYWVVGRLFLHPKPQELYVCFHDSVTEIAIEIAFKLFFGLTL

>FUZZY
MGEEGTGGTVHLLCLAASSGVPLFCRSSRGGAPARQQLPFSVIGSLNGVHMFGQNLEVQLSSARTENTTVVWKSFHDSITLIVLSSEVGISELRLERLLQMVFGAMVLLVGLEELTNIRNVERLKKDLRASYCLIDS
FLGDSELIGDLTQCVDCVIPPEGSLLQEALSGFAEAAGTTFVSLVVSGRVVAATEGWWRLGTPEAVLLPWLVGSLPPQTARDYPVYLPHGSPTVPHRLLTLTLLPSLELCLLCGPSPPLSQLYPQLLERWWQPLL
DPLRACLPLGPRALPSGFPLHTDILGLLLLHLELKRCLFTVEPLGDKEPSPEQRRRLLRNFYTLVTSTHFPPEPGPPEKTEDEVYQAQLPRACYLVLGTEEPGTGVRLVALQLGLRRLLLLLSPQSPTHGLRSLATH
TLHALTPLL

>RSG1
MARPPVPGSVVVPNWHESAEGKEYLACILRKNRRRVFGLLERPVLLPPVSIDTASYKIFVSGKSGVGKTALVAKLAGLEVPVVHHETTGIQTTVVFWPAKLQASSRVVMFRFEFWDCGESALKKFDHMLLACMEN
TDAFLFLFSFTDRASFEDLPGQLARIAGEAPGVVRMVIGSKFDQYMHTDVPERDLTAFRQAWELPLLRVKSVPGRRLADGRTLDGRAGLADVAHILNGLAEQLWHQDQVAAGLLPNPPESAPE

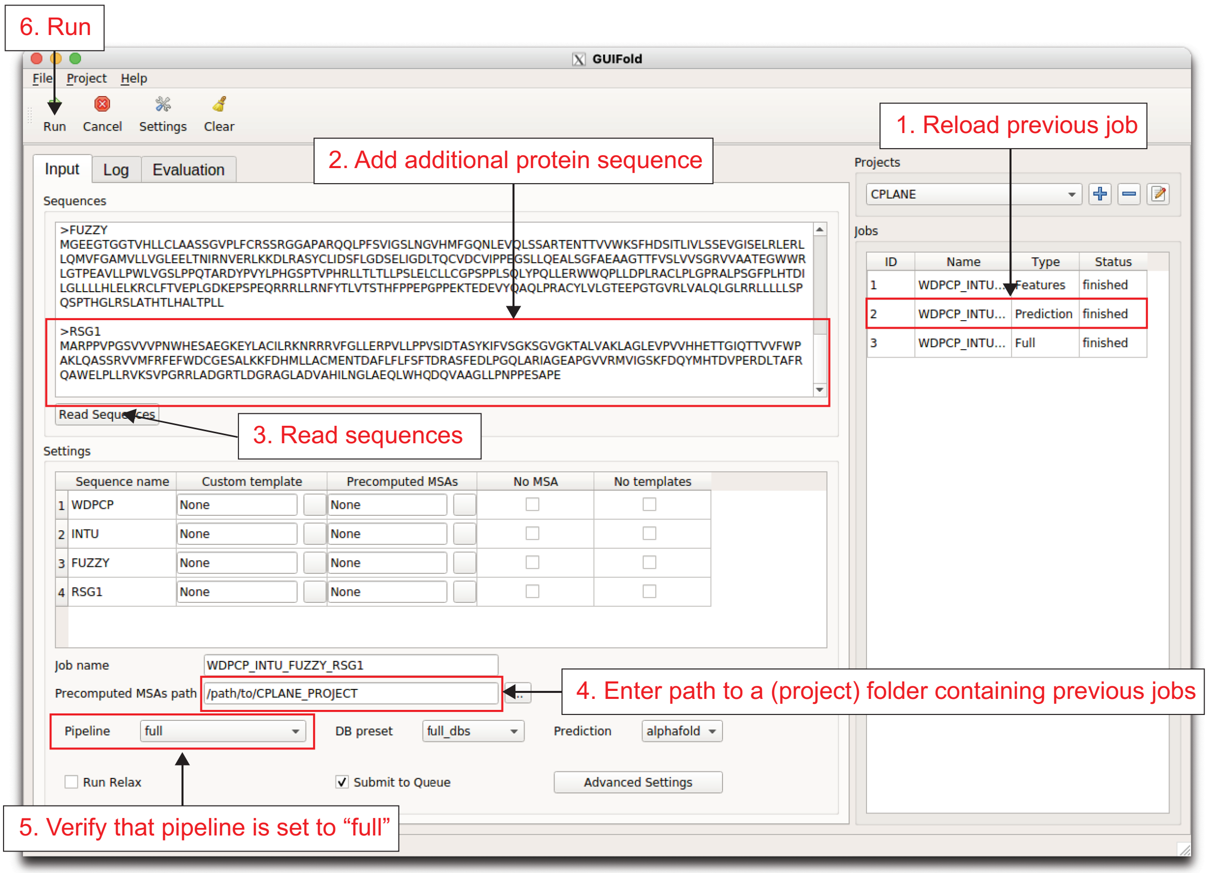


**Supplemental Figure 4: Screenshot of input when re-using previous MSAs.** This example shows how to predict the structure of the CPLANE-RSG1 complex. The MSAs from the previous job (CPLANE core complex) can be re-used and only the MSAs for RSG1 need to be calculated. When giving the project folder as precomputed MSAs path, the pipeline will search all subfolders for matching MSAs. Alternatively, for each component, the path to the respective folder containing matching MSA files can be given in the component table.

#### 1.6 Full evaluation results for CPLANE-RSG1 prediction

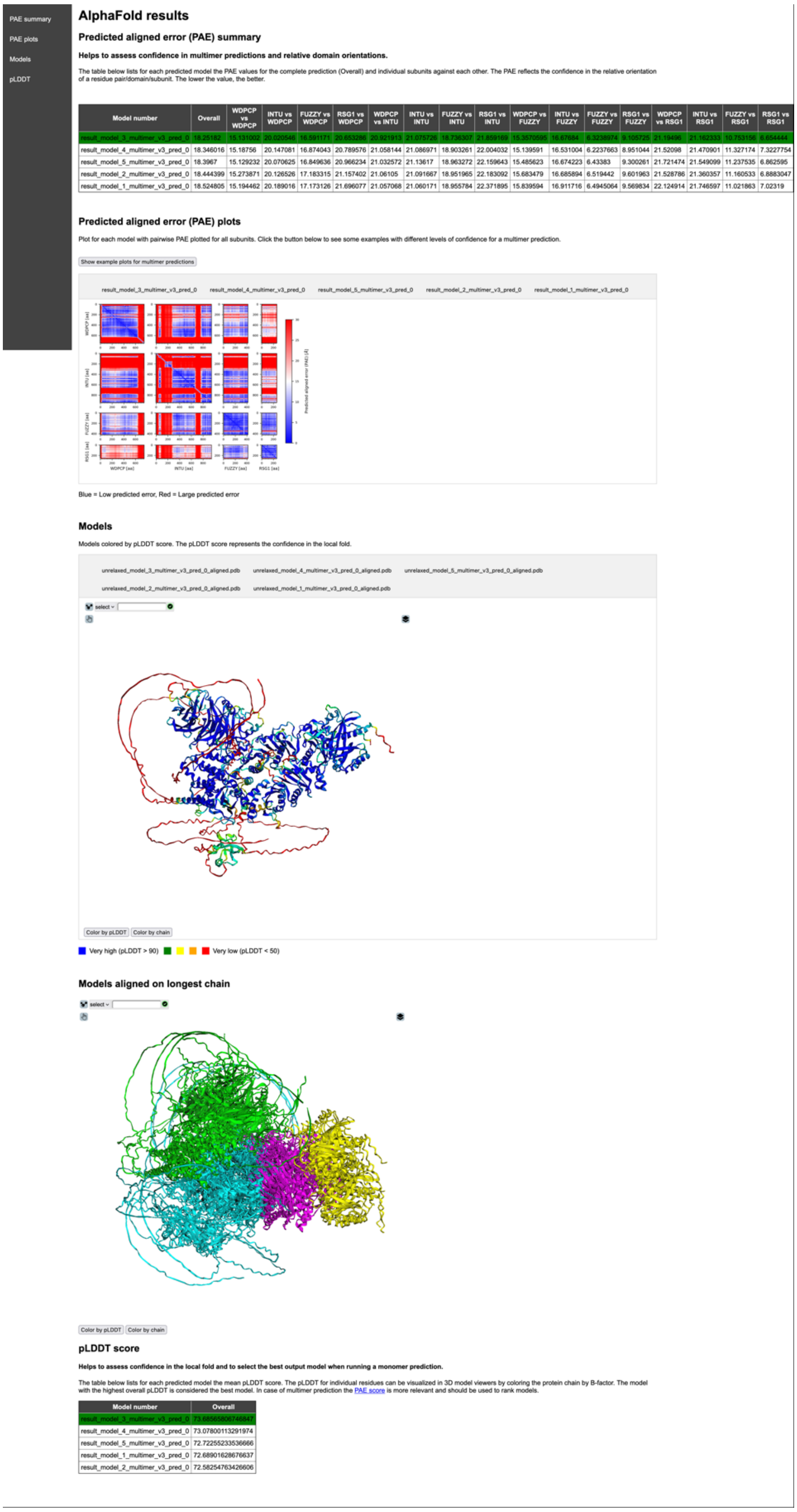

PAE summary
PAE plots
Models
pLDDT

##### AlphaFold results

###### Predicted aligned error (PAE) summary

**Helps to assess confidence in multimer predictions and relative domain orientations.**

The table below lists for each predicted model the PAE values for the complete prediction (Overall) and individual subunits against each other. The PAE reflects the confidence in the relative orientation of a residue pair/domain/subunit. The lower the value, the better.

| Model number | Overall | WDPCP vs WDPCP | INTU vs WDPCP | FUZZY vs WDPCP | RSG1 vs WDPCP | WDPCP vs INTU | INTU vs INTU | FUZZY vs INTU | RSG1 vs INTU | WDPCP vs FUZZY | INTU vs FUZZY | FUZZY vs FUZZY | RSG1 vs FUZZY | WDPCP vs RSG1 | INTU vs RSG1 | FUZZY vs RSG1 | RSG1 vs RSG1 |
|---|---|---|---|---|---|---|---|---|---|---|---|---|---|---|---|---|---|
| result_model_3_multimer_v3_pred_0 | 18.25182 | 15.131002 | 20.020546 | 16.591171 | 20.653286 | 20.921913 | 21.075726 | 18.736307 | 21.859169 | 15.3570595 | 16.67684 | 6.3238974 | 9.105725 | 21.19496 | 21.162333 | 10.753156 | 6.654444 |
| result_model_4_multimer_v3_pred_0 | 18.346016 | 15.18756 | 20.147081 | 16.874043 | 20.789576 | 21.058144 | 21.086971 | 18.903261 | 22.004032 | 15.139591 | 16.531004 | 6.2237663 | 8.951044 | 21.52098 | 21.470901 | 11.327174 | 7.3227754 |
| result_model_5_multimer_v3_pred_0 | 18.3967 | 15.129232 | 20.070625 | 16.849636 | 20.966234 | 21.032572 | 21.13617 | 18.963272 | 22.159643 | 15.485623 | 16.674223 | 6.43383 | 9.300261 | 21.721474 | 21.549099 | 11.237535 | 6.862595 |
| result_model_2_multimer_v3_pred_0 | 18.444399 | 15.273871 | 20.126526 | 17.183315 | 21.157402 | 21.06105 | 21.091667 | 18.951965 | 22.183092 | 15.683479 | 16.685894 | 6.519442 | 9.601963 | 21.528786 | 21.360357 | 11.160533 | 6.8883047 |
| result_model_1_multimer_v3_pred_0 | 18.524805 | 15.194462 | 20.189016 | 17.173126 | 21.696077 | 21.057068 | 21.060171 | 18.955784 | 22.371895 | 15.839594 | 16.911716 | 6.4945064 | 9.569834 | 22.124914 | 21.746597 | 11.021863 | 7.02319 |

###### Predicted aligned error (PAE) plots

Plot for each model with pairwise PAE plotted for all subunits. Click the button below to see some examples with different levels of confidence for a multimer prediction.

Show example plots for multimer predictions

result_model_3_multimer_v3_pred_0 result_model_4_multimer_v3_pred_0 result_model_5_multimer_v3_pred_0 result_model_2_multimer_v3_pred_0 result_model_1_multimer_v3_pred_0



Blue = Low predicted error, Red = Large predicted error

###### Models

Models colored by pLDDT score. The pLDDT score represents the confidence in the local fold.

unrelaxed_model_3_multimer_v3_pred_0_aligned.pdb unrelaxed_model_4_multimer_v3_pred_0_aligned.pdb unrelaxed_model_5_multimer_v3_pred_0_aligned.pdb
unrelaxed_model_2_multimer_v3_pred_0_aligned.pdb unrelaxed_model_1_multimer_v3_pred_0_aligned.pdb

select

Color by pLDDT Color by chain

Very high (pLDDT > 90) Very low (pLDDT < 50)

###### Models aligned on longest chain

select

Color by pLDDT Color by chain

###### pLDDT score

**Helps to assess confidence in the local fold and to select the best output model when running a monomer prediction.**

The table below lists for each predicted model the mean pLDDT score. The pLDDT for individual residues can be visualized in 3D model viewers by coloring the protein chain by B-factor. The model with the highest overall pLDDT is considered the best model. In case of multimer prediction the PAE score is more relevant and should be used to rank models.

| Model number | Overall |
|---|---|
| result_model_3_multimer_v3_pred_0 | 73.68565806746847 |
| result_model_4_multimer_v3_pred_0 | 73.07800113291974 |
| result_model_5_multimer_v3_pred_0 | 72.72255233536666 |
| result_model_1_multimer_v3_pred_0 | 72.68901628676637 |
| result_model_2_multimer_v3_pred_0 | 72.58254763426606 |

**Supplemental Figure 5: Screenshot of evaluation results (CPLANE-RSG1 prediction).**
